# Estimating Drivers of Cell State Transitions Using Gene Regulatory Network Models

**DOI:** 10.1101/089003

**Authors:** Daniel Schlauch, Kimberly Glass, Craig P. Hersh, Edwin K. Silverman, John Quackenbush

## Abstract

Specific cellular states are often associated with distinct gene expression patterns. These states are plastic, changing during development, or in the transition from health to disease. One relatively simple extension of this concept is to recognize that we can classify different cell-types by their active gene regulatory networks and that, consequently, transitions between cellular states can be modeled by changes in these underlying regulatory networks. Here we describe **MONSTER**, **MO**deling **N**etwork **S**tate **T**ransitions from **E**xpression and **R**egulatory data, a regression-based method for inferring transcription factor drivers of cell state conditions at the gene regulatory network level. As a demonstration, we apply MONSTER to four different studies of chronic obstructive pulmonary disease to identify transcription factors that alter the network structure as the cell state progresses toward the disease-state. Our results demonstrate that MONSTER can find strong regulatory signals that persist across studies and tissues of the same disease and that are not detectable using conventional analysis methods based on differential expression. An R package implementing MONSTER is available at github.com/QuackenbushLab/MONSTER.

## Introduction

Cell state phenotypic transitions, such as those that occur during development, or as healthy tissue transforms into a disease phenotype, are fundamental processes that operate within biological systems. Understanding what drives these transitions, and modeling the processes, is one of the great open challenges in modern biology. One way to conceptualize the state transition problem is to imagine that each phenotype has its own characteristic gene regulatory network, and that there are a set of processes that are either activated or inactivated to transform the network in the initial state into one that characterizes the final state. Identifying those changes could, in principle, help us to understand not only the processes that drive the state change, but also how one might intervene to either promote or inhibit such a transition.

Each distinct cell state consists of a set of characteristic processes, some of which are shared across many cell-states (“housekeeping” functions) and others which are unique to that particular state. These processes are controlled by gene regulatory networks in which transcription factors (and other regulators) moderate the transcription of individual genes whose expression levels, in turn, characterize the state. One can represent these regulatory processes as a directed network graph, in which transcription factors and genes are nodes in the network, and edges represent the regulatory interactions between transcription factors and their target genes. A compact representation of such a network, with interactions between *m* transcription factors and *p* target genes, is as a binary *p* × *m* “adjacency matrix”. In this matrix, a value of 1 represents an active interaction between a transcription factor and a potential target, and 0 represents the lack of a regulatory interaction.

When considering networks, a cell state transition is one that transforms the initial state network to the final state network, adding and deleting edges as appropriate. Using the adjacency matrix formalism, one can think of this as a problem in linear algebra in which we attempt to find an *m* × *m* “transition matrix” **T**, subject to a set of constraints, that approximates the conversion of the initial network's adjacency matrix **A** into the final network's adjacency matrix **B**, or

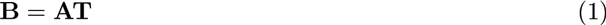

In this model, the diagonal elements of **T** map network edges to themselves. The drivers of the transition are those off-diagonal elements that change the configuration of the network between states.

While this framework, as depicted in Figure 1, is intuitive, it is a bit simplistic in that we have cast the initial and final states as discrete. However, the model can be generalized by recognizing that any phenotype we analyze consists of a collection of individuals, all of whom have a slightly different manifestation of the state, and therefore a slightly different active gene regulatory network. Practically, what that means is that for each state, rather than having a network model with edges that are either “on” or “off,” a phenotype should be represented by a network in which each edge has a weight that represents an estimation of its presence across the population. In other words, the initial and final state adjacency matrices are not comprised of 1's and 0's, but of continuous variables that estimate population-level regulatory network edge-weights. Consequently, the problem of calculating the transition matrix is generalized to solving **B** = **AT** + **E**, where **E** is an *p* × *m* error matrix. In this expanded framework, modeling the cell state transition remains equivalent to estimating the appropriate transition matrix **T**, and then identifying state transition drivers based on features of that matrix.

**Figure 1:**
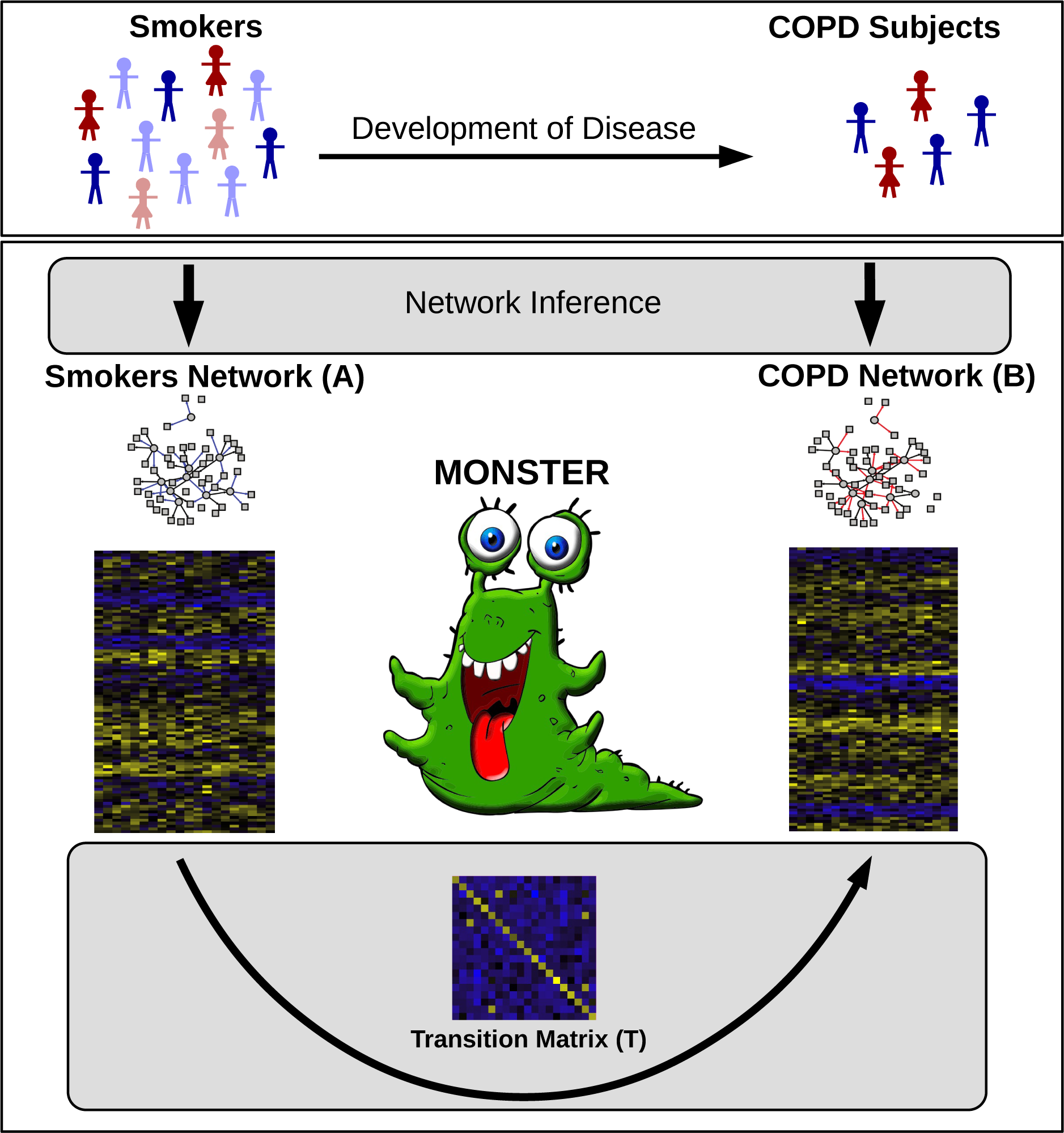
Overview of the MONSTER approach, as applied to the transition between smokers and those suffering from chronic obstructive pulmonary disease (COPD). MONSTER's approach seeks to find the *T F T F* transition matrix that best characterizes the state change in network structure between the initial and final biological conditions. Subjects are first divided into two groups based on whether they have COPD or are smokers that have not yet developed clinical COPD. Network inference is then performed separately on each group, yielding a bipartite adjacency matrix connecting transcription factors to genes. Finally, a transition matrix is computed which characterizes the conversion from the consensus Smokers Network to the COPD Network.

## Significance Statement

Biological states are characterized by distinct patterns of gene expression that reflect each phenotype's active cellular processes. Driving these phenotypes are gene regulatory networks in which transcriptions factors control when and to what degree individual genes are expressed. Phenotypic transitions, such as those that occur when disease arises from healthy tissue, are associated with changes in these networks. MONSTER is a new approach to understanding these transitions. MONSTER models phenotypic-specific regulatory networks and then estimates a “transition matrix” that converts one state to another. By examining the properties of the transition matrix, we can gain insight into regulatory changes associated with phenotypic state transition. We demonstrate the power of MONSTER by applying it to data from four independent studies of chronic obstructive pulmonary disease and find a robust set of transcription factors that help explain the development of the disease.

## MONSTER: MOdeling Network State Transitions from Expression and Regulatory data

The MONSTER algorithm models the regulatory transition between two cellular states in three steps: (1) Inferring state-specific gene regulatory networks, (2) modeling the state transition matrix, and (3) computing the transcription factor involvement.

**Inferring state-specific gene regulatory networks**: Before estimating the transition matrix, T, we must first estimate a gene regulatory starting point for each state. While there have been many methods developed to infer such networks [24, 19, 20, 11, 4, 36, 44], we have found the bipartite framework used in PANDA [18] to have features that are particularly amenable to interpretation in the context of state transitions. PANDA begins by using genome-wide transcription factor binding data to postulate a network “prior”, and then uses message-passing to integrate multiple data sources, including state-specific gene co-expression data.

Motivated by PANDA, we developed a highly computationally efficient, classification-based network inference method that uses common patterns between transcription factor targets and gene co-expression to estimate edges and to generate a bipartite gene regulatory network connecting transcription factors to their target genes.

This approach is based on the simple concept that genes affected by a common transcription factor are likely to exhibit correlated patterns of expression. To begin, we combine gene co-expression information with information about transcription factor targeting derived from sources such as ChIP-Seq or sets of known sequence binding motifs found in the vicinity of genes. we then calculate the direct evidence for a regulatory interaction between a transcription factor and gene, which we define as the squared partial correlation between a given transcriptionfactor's gene expression, *g_i_*, and the gene's expression, *g_j_*, conditional on all other trnscriptionfactors’gene expression:

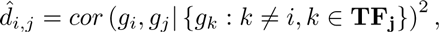

where *g_i_* is the gene which encodes the transcription factor *TF_i_*, *g_j_* is any other gene in the genome, and **TF_j_** is the set of gene indices corresponding to known transcription factors with binding site in the promoter region of *g_j_*. The correlation is conditioned on the expression of all other potential regulators of *g_j_* based on the transcription factor motifs associated with *g_j_*.

Next, we fit a logistic regression model which estimates the probability of each gene, indexed *j*, being a motif target of a transcription factor, indexed *i*, based on the expression pattern across the *n* samples across *p* genes in each phenotypic class:

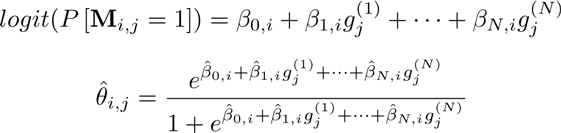

where the response **M** is a binary *p* × *m* matrix indicating the presence of a sequence motif for the *i^th^* transcription factor in the vicinity of each of the *j^th^* gene. And where 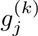 represents the gene expression measured for sample *k* at gene *j*. Thus, the fitted probability *θ̂_i,j_* represents our estimated indirect evidence. Combining the scores for the direct evidence, *d̂_i,j_*, and indirect evidence, *θ̂_i,j_*, via weighted sum between each transcription factor-gene pair yields estimated edge-weights for the gene regulatory network (see Supporting Information).

Applying this approach to gene expression data from two distinct phenotypes results in two *p* × *m* gene regulatory adjacency matrices, one for each phenotype. These matrices represent estimates of the targeting patterns of the *m* transcription factors onto the *p* genes. This network inference algorithm finds validated regulatory interactions in Escherichia coli and Yeast (Saccharomyces cerevisiae) data sets (see Supporting Information).

**Modeling the state transition matrix**: Once we have gene regulatory network estimates for each phenotype, we can formulate the problem of estimating the transition matrix in a regression framework in which we solve for the *m* × *m* matrix that best describes the transformation between phenotypes (1). More specifically, MONSTER predicts the change in edge-weights for a transcription factor, indexed *i*, in a network based on all of the edge-weights in the baseline phenotype network.

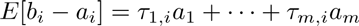

where *b_i_* and *a_i_* are column-vectorsin **B** and **A** thatdescribethe regulatorytargetingof transcriptionfactor*i* in the final and initialnetworks,respectively.

In the simplest case, this can be solved with normal equations,

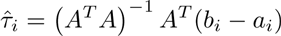

to generateeach of the columns of the transitionmatrix **T** such that

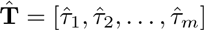

The regression is performed *m* times corresponding to each of the transcription factors in the data. In this sense, columns in the transition matrix can be loosely interpreted as the optimal linear combination of columns in the initial state adjacency matrix which predict the column in the final state adjacency matrix. (see Supporting Information).

This framework allows for the natural extension of constraints such as *L*1 and/or *L*2 regularization (see Supporting Information). For the analysis we present in this manuscript, we use the normal equations and do not impose a penalty on the regression coefficients.

**Computing the transcription factor involvement**: For a transition between two nearly identical states, we expect that the transition matrix would approximate the identity matrix. However, as initial and final states diverge, there should be increasing differences in their corresponding gene regulatory networks and, consequently, the transition matrix will also increasingly diverge from the identity matrix. In this model, the transcription factors that most significantly alter their regulatory targets will have the greatest “off-diagonal mass” in the transition matrix, meaning that they will have very different targets between states and so are likely to be involved in the state transition process. We define the “differential transcription factor involvement” (dTFI) as the magnitude of the off-diagonal mass associated with each transcription factor, or,

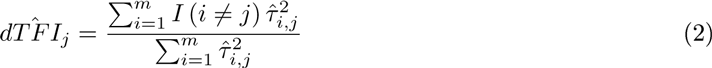

where, 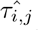 is the value in of the element *i^th^* row and *j^th^* column in the transition matrix, corresponding to the *i^th^* and *j^th^* transcription factors. To estimate the significance of this statistic, we randomly permute sample labels *n* = 400 times across phenotypes (see Supporting Information).

## MONSTER finds significantly differentially involved transcription factors in COPD with strong concordance in independent data sets

As a demonstration of the power of MONSTER to identify driving factors in disease, we applied the method to case-control gene expression data sets from four independent Chronic Obstructive Pulmonary Disease (COPD) cohorts: Evaluation of COPD Longitudinally to Identify Predictive Surrogate End-points (ECLIPSE) [45][49] (2), COPDGene [42] [2] [37], Lung Genomics Research Consortium (LGRC) [1] and Lung Tissue from Channing Division of Network Medicine (LT-CDNM) [39]. The tissues assayed in ECLIPSE and COPDGene were whole blood and peripheral blood mononuclear cells (PBMCs), respectively, while homogenized lung tissue was sampled for LGRC and LT-CDNM.

As a baseline comparison metric, we evaluated the efficacy of applying commonly used network inference methods on these case-control studies. In analyzing phenotypic changes, networks are generally compared directly, with changes in the presence or weight of edges between key genes being of primary interest. It is therefore reasonable to assume that any reliable network results generated from a comparison of disease to controls will be reproducible in independent studies. We investigated whether this is the case for our four COPD data sets using three widely used network inference methods - Algorithm for the Reconstruction of Gene Regulatory Networks (ARACNE)[34], Context Likelihood of Relatedness (CLR)[13], and Weighted Gene Correlation Network Analysis (WGCNA) [50] - computing the difference in edge weights between cases and controls for each of the four studies. We found no meaningful correlation (*R*^2^ < .01) of edge weight difference across any of the studies regardless of network inference method or tissue type (Supporting Figure 3). Edge weight differences, even when very large in one study, did not reproduce in other studies. This suggests that a simple direct comparison of edges between inferred networks is insufficient for extracting reproducible drivers of network state transitions. This finding may be unsurprising given the difficulty in inferring individual edges in the presence of heterogeneous phenotypic states, technical and biological noise with a limited number of samples.

The lack of replication in edge-weight differences between independent data sets representing similar study designs indicates that we need to rethink how we evaluate network state transitions. MONSTER provides a unique approach for making that comparison. In each of the four COPD data sets, we used MONSTER to calculate the differential transcription factor involvement (*dTFI*, Equation 2) for each transcription factor and used permutation analysis to estimate their significance (Figure 2, Additional Figures 1-3). We observed strongly significant (*p* < 1*e* 15) correlation in dTFI values for each pairwise combination of studies. In addition, out of the top 10 most differentially involved transcription factors in the ECLIPSE and COPDGene studies, we found 7 to be in common. Furthermore, three of these seven transcription factors (GABPA, ELK4, ELK1) also appeared as significant in the LGRC results with FDR<0.01 and each of the top five ECLIPSE results were among the top seven in the LT-CDNM results (Additional Table 1, Additional Figure 3). This agreement is quite striking considering that the there was almost no correlation in the edge-weight differences across these same studies when we tested the other methods. But it is exactly what we should expect-that the same method applied to independent studies of the same phenotypes should produce largely consistent results.

**Figure 2:**
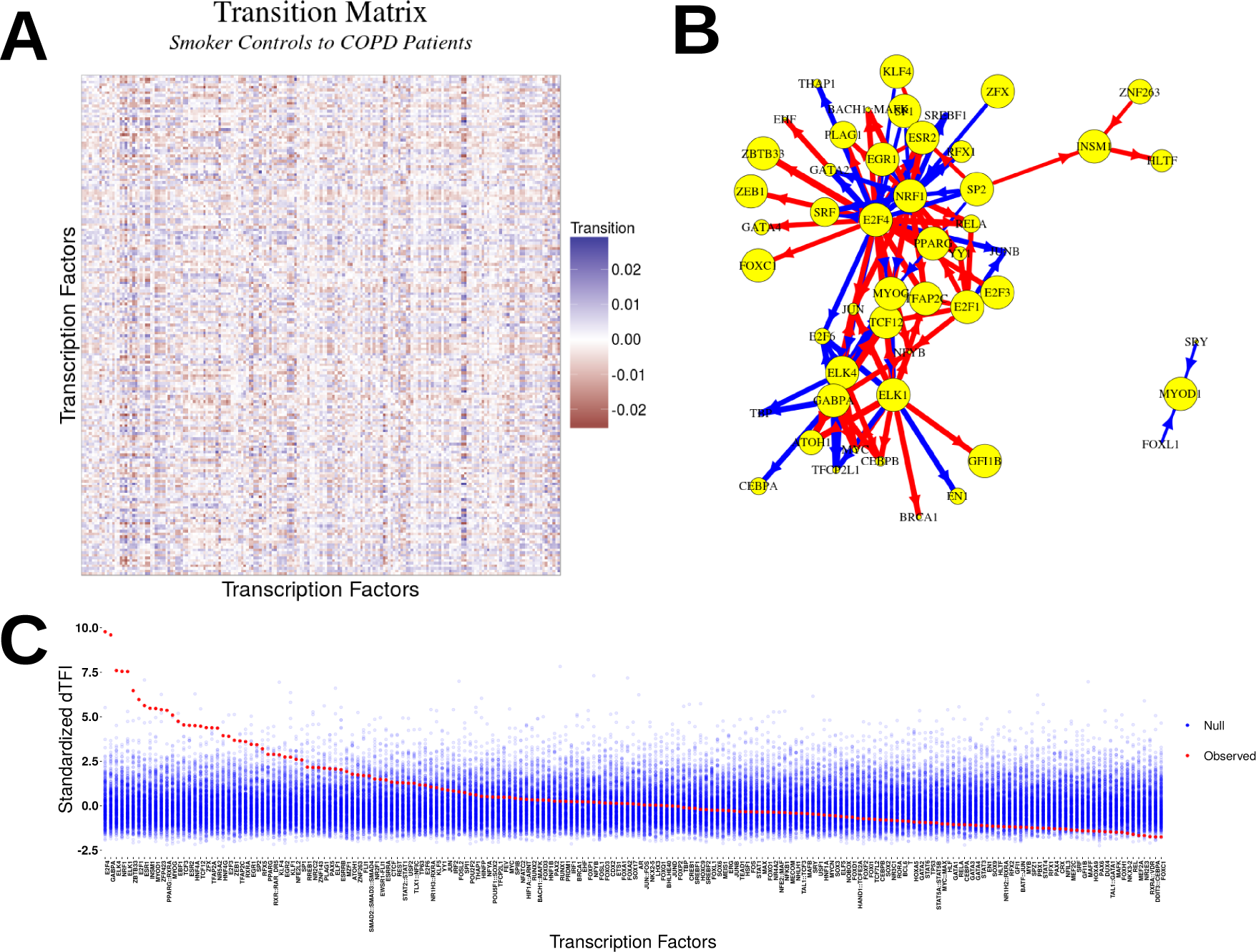
MONSTER analysis results in the ECLIPSE study. **A** Heatmap depicting the transition matrix calculated for smoker controls “transitioning” to COPD by applying MONSTER to ECLIPSE gene expression data. For the purposes of visualization, the magnitude of the diagonal is set to zero. **B** A network visualization of the 100 largest transitions identified based on the transition matrix in (A). Arrows indicate a change in edges from a transcription factor in the Smoker-Control network to resemble those of a transcription factor in the COPD network. Edge thickness represents the magnitude of the transition and node (TFs) sizes represent the dTFI for that TF. Blue edges represent a gain of targeting features and red represents the loss. **C** The dTFI score from MONSTER (red) and the background null distribution of dTFI values (blue) as estimated by 400 random sample permutations of the data.

Many of the top dTFI transcription factors, especially those identified by MONSTER across all four studies, are biologically plausible candidates to be involved in the etiology of COPD (Additional Table 1, Additional Figures 1-3). For example, E2F4 is a transcriptional repressor important in airway development [9] and studies have begun to demonstrate the relevance of developmental pathways in COPD pathogenesis [3].

Some of the greatest effect sizes across all four studies were found for SP1 and SP2. An additional member of the SP transcription factor family, SP3, has been shown to regulate HHIP, a known COPD susceptibility gene [51]. Both SP1 and SP2 form complexes with the E2F family [43, 28] and may play a key role in the alteration of E2F4 targeting behavior. Furthermore, E2F4 has been found to form a complex with EGR-1 (a highly significant transcription factor in ECLIPSE and LT-CDNM) in response smoke exposure, which may lead to autophagy, apoptosis and subsequently to development of emphysema [5].

Mitochondrial mechanisms have also been associated with COPD progression [7]. Two of most highly significant transcription factors based on dTFI in ECLIPSE were NRF1 and GABPA (FDR<.001). Indeed, these TFs had highly significant dTFI (FDR<0.1) in all four studies. NRF1 regulates the expression of nuclear encoded mitochondrial proteins [21]. GABPA, also known as human nuclear respiratory factor‐ 2 subunit alpha, may have a similar role in nuclear control of mitochondrial gene expression. Furthermore, GABPA interacts with SP1 [16] providing evidence of a potentially shared regulatory mechanism with E2F4.

Overall, we found a strong correlation across studies in transcription factors identified as significantly differentially involved (Figure 3A-3B). It is reassuring that we find the strongest agreement when comparing studies that assayed similar tissues. However the fact that we see similar dTFI signal across studies involving different tissue types is also notable as it suggests that the transition from smoker control to disease phenotype affects multiple tissues and supports the growing evidence for a role in immune response in COPD pathogenesis.

**Figure 3:**
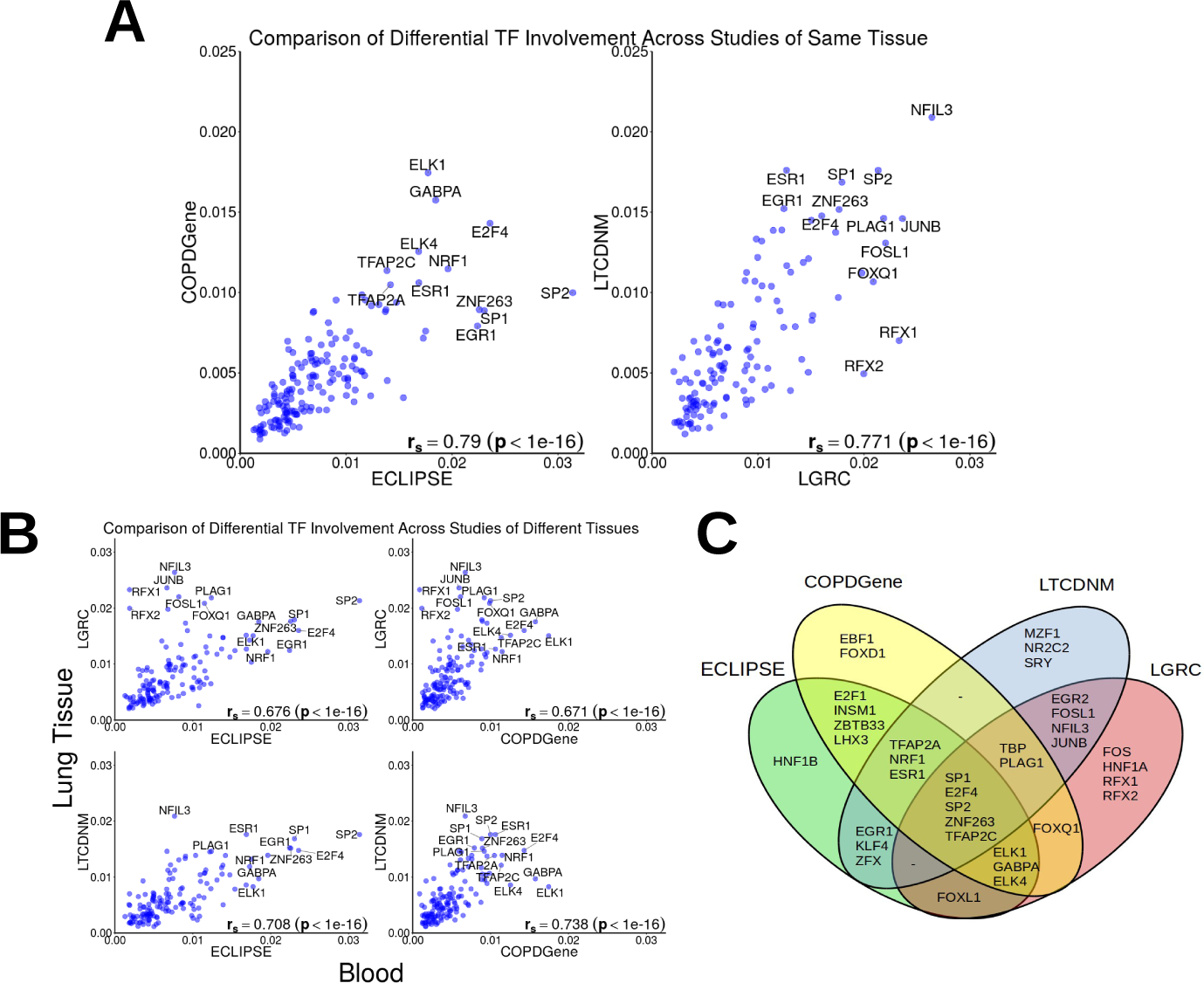
Strong reproducibility in top differential transcription factor involvement found in case-control COPD studies. ECLIPSE and COPDGene profiled gene expression in whole-blood and PBMC while the gene expression data in LGRC and LT-CDNM were assayed in lung tissue. **A** Results for studies with gene expression data obtained from the same-tissue. Both the blood based (left) and lung tissue studies (right) demonstrate very high Spearman correlation of differential involvement. **B** Despite using data from different sources we found agreement between studies of different tissues. **C** Venn diagram depicting the top 20 transcription factors found in each study. The union of all top 20 lists contains 36 transcription factors.

Gene regulatory networks, and results derived from their comparison, are notoriously difficult to replicate across studies[46]. The four studies we used each has unique aspects, including the choice of microarray platform, study demographics, location, time, and tissue. Nevertheless, MONSTER identified similar sets of transcription factors associated with the transition between cases and controls. This consistency in biologically-relevant transcription factors, associated with the transition from the control phenotype to disease, in four independent studies suggests that MONSTER can provide not only robust network models, but also can identify reliable differences between networks.

Despite the overall consistency, some transcription factors had variable *dTFI* across studies. For example, using the LGRC dataset, we discovered a highly significant (*F DR* < .0001) differential targeting pattern involving the transcription factors RFX1 and RFX2 (Additional Table 1). However, these same TFs were not identified as potential drivers of the control to COPD transition in either the ECLIPSE or COPDGene study. This difference is likely due the differences in tissue type as the RFX family transcription factors are known to regulate ciliogenesis [6]. Cilia are critical for clearing mucous from the airways of healthy individuals, but disruption can lead to infection and potentially to chronic airflow obstruction [23, 25, 12].

The hypothesis behind MONSTER is that each phenotype has a unique gene regulatory network and that a change in phenotypic state is reflected in changes in transcription factor targeting. That hypothesis translates to an expectation that transcription factors driving change in phenotype will have the greatest *dTFI* scores. One might expect that these “driving transcription” factors would also be differentially expressed. We compared *dTFI* to differential expression (ECLIPSE Figure 4, other studies shown in Additional Figure 4) and found that many of the transcription factors with high dTFI values were not differentially expressed. This suggests that there are other mechanisms, such as epigenetic modification of the genome or protein modifications, that alter the structure of the regulatory network by changing which genes are targeted by key transcription factors.

**Figure 4:**
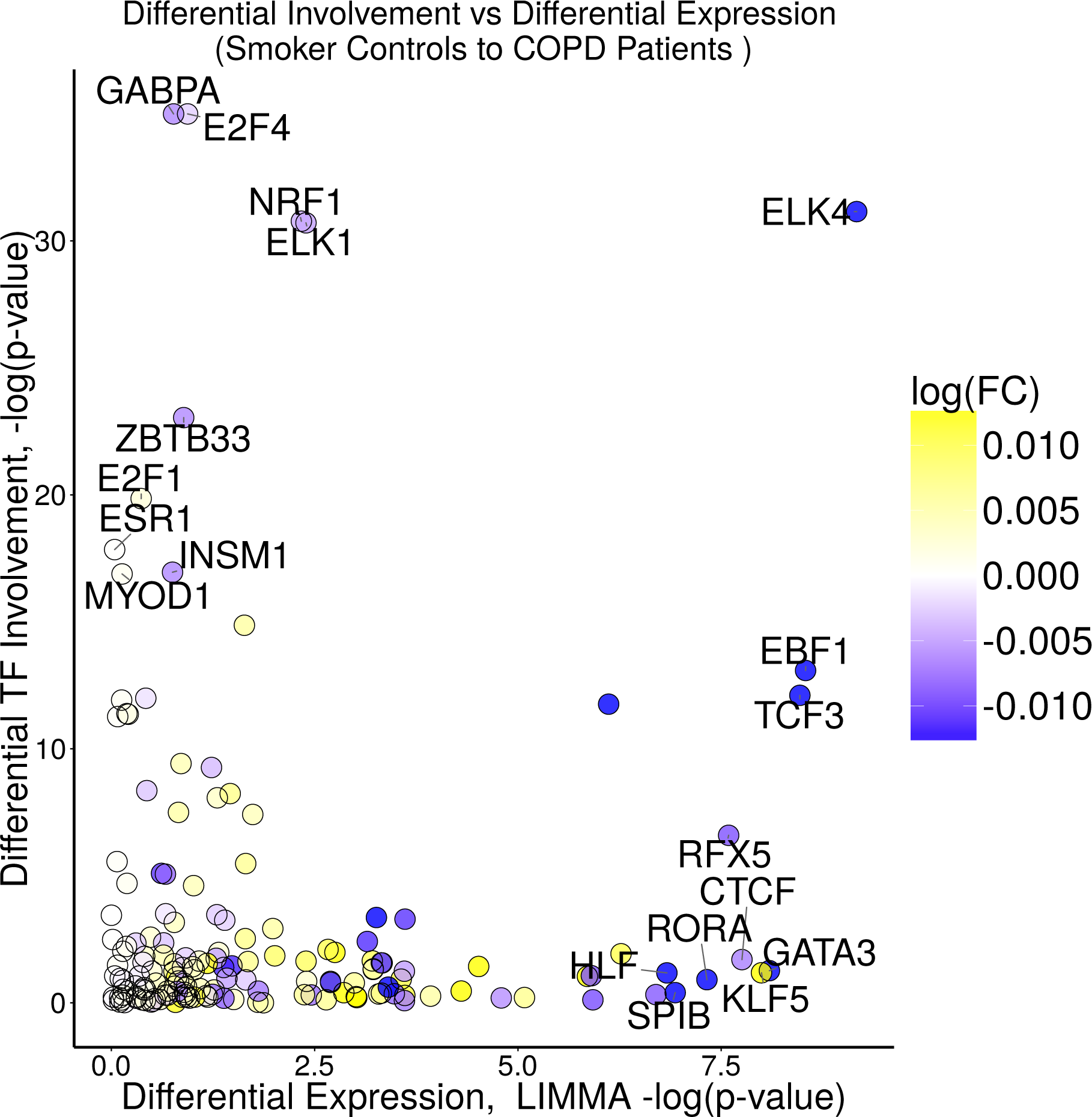
Differentially involved transcription factors are not necessarily differentially expressed. A plot of the differential expression versus the differential involvement for transcription factors based on our analysis of the ECLIPSE data. MONSTER commonly finds transcription factors which are differentially involved but are expressed at similar levels across cases and controls. Importantly, these transcription factors would not have been identified using conventional differential expression methods. This demonstrates the unique potential MONSTER has for discovery beyond standard gene expression analysis.

## Discussion

One of the fundamental problems is biology is modeling the transition between biological states such as that which occurs during development or as a healthy tissue transforms into a disease state. As our ability to generate large-scale, integrative multi-omic data sets has grown, there has been an increased interest in using those data to infer gene regulatory networks to model fundamental biological processes. There have been many network inference methods published, each of which uses a different approach to estimating the “strength” of interactions between genes (or between transcription factors and their targets). But all suffer from the same fundamental limitation: every method relies on estimating weights that represent the likelihood of an interaction between two genes to identify “real” (high confidence) edges. In comparing phenotypes, most methods then subtract discretized edges in one phenotype from those in the other to search for differences.

MONSTER represents a new way of looking at phenotypic transitions, but one that captures many aspects of what we should expect. First, we have to recognize that there is no single network that represents a phenotype, but that each phenotype is represented by a family of networks that all vary slightly from each other, yet which have essential features that are consistent with the phenotype. What this means is that each regulatory edge in a network representation has to be represented by continuous, rather than discrete, variables. This captures the fact that regulatory interactions are stronger in certain individuals and weaker in others, or present in some and absent in others, but that, on average, they represent a distribution.

Second, when we consider a change in phenotype, that will be reflected in altered patterns of gene expression, and ultimately in the networks that represent the phenotype. In a transition, some individuals will experience a greater change while others will experience a smaller change. But overall, regulatory patterns in the network will shift as the phenotype changes.

Third, the change in the gene regulatory network structure between phenotypes will be driven by changes in the connectivity of the regulators-the transcription factors that alter when, how, and how strongly genes are expressed. A natural hypothesis in this model is that the transition between phenotype is likely associated with the transcription factors that experience the greatest change in their regulatory patterns between states, and that the activation or inactivation of their target genes, and the functions carried out by those genes, likely reflect the phenotypic differences between states.

MONSTER captures these features, creating initial and final state network representations and estimating the change in transcription factor regulatory patterns by estimating a transition matrix. For each transcription factor, the “off diagonal mass” calculated as the differential transcription factor involvement (dTFI), identifies those transcription factors that are ultimately likely to drive the phenotypic state transition.

In applying MONSTER to four independent COPD gene expression data sets surveying both COPD and smoker controls, a highly consistent picture of the transcription factors associated with disease development emerges. This consistency is, to some, surprising as gene expression data is notoriously noisy, with each study finding sets of differentially expressed genes that often are not concordant. By focusing on transcriptional regulators, MONSTER seems to be able to separate a cleaner signal from the noise and one that makes some biological sense. Indeed, when one looks at the transcription factors found by MONSTER as associated with the transition, all are biologically plausible candidates which provide new and important opportunities for future molecular studies of COPD pathogenesis. It is also noteworthy that many of these transcription factors could not have been found through a simple differential expression analysis as their transcriptional levels do not change significantly between disease and control populations. Rather, it is the regulatory patterns of these transcription factors, possibly driven by epigenetic or other changes, that shifts with the phenotype.

## Acknowledgements

The project described was supported by Award Number P01 HL105339, R01 HL111759, R01 HL089897, R01 HL089856 and K25HL133599 from the National Heart, Lung, and Blood Institute. The content is solely the responsibility of the authors and does not necessarily represent the official views of the National Heart, Lung, and Blood Institute or the National Institutes of Health.

## Author Contributions

DS, KG, and JQ designed research; DS performed method development and application; DS, KG, CPH, EKS and JQ interpreted results; DS, KG, CPH, EKS and JQ wrote the paper. The authors declare no conflict of interest. 5 To whom correspondence should be addressed. E-mail: johnqjimmy.harvard.edu

## Overview of Supporting Information

This Supporting Information document is broken into four main sections that include:

- A description of the data used for the COPD network inference and analysis presented in the main text
- A detailed description of the MONSTER approach for defining network state transitions
- Various evaluations of the MONSTER method
- An illustration of the irreproducibility of network differences outside of the transition matrix formalism

## Data for COPD Network Inference and Analysis

### Sequence binding motifs

A regulatory network prior between transcription factors and target genes was created by using position weight matrices for 205 transcription factor motifs obtained from JASPAR 2014 (http://jaspar2014.genereg.net/), [35] and running Haystack[38] to scan the hg19 genome for occurrences of these motifs. Sequences were identified as hits for a transcription factor if they satisfied the significance threshold of *p* < 10^−5^. We then used HOMER (http://homer.salk.edu/homer/ngs/index.html) [22] to identify transcription factor binding motifs that map to a window ranging from 750 base pairs downstream to 250 base pairs upstream of each gene's transcription start site under the assumption that transcription factors falling in this region may actively regulate expression of the gene.

### ECLIPSE

Gene expression data from the ECLIPSE study (GSE54837) [45] was collected using blood samples from 226 subjects classified as non-smokers (6), smoker controls (84) or COPD (136). Blood samples from each individual were profiled using Affymetrix Human Genome U133 Plus 2.0 microarrays. CEL data files from these assays were RMA-normalized[26] in R using the Bioconductor package ‘affy’[17]. Array probes were collapsed to 19,765 Entrez-gene IDs using a custom CDF[8] and the 220 samples for COPD or smoker control subjects were retained for analysis. Finally, genes were associated with potential regulatory transcription factors using a motif scan (described above). 1,553 genes were not associated with any transcription factor and excluded from further analysis, leaving 17,342 genes that were used to construct network models.

### COPDGene

Gene expression data from the COPDGene study (GSE42057) [2, 42] was collected from blood samples obtained from 136 subjects classified as smoker controls (42) or COPD (94) and profiled on Affymetrix Human Genome U133 Plus 2.0 microarrays. Similar to the ECLIPSE data, CEL data files from these microarray assays were RMA-normalized using the ‘affy’ package and array probes were collapsed to Entrez-gene IDs using a custom CDF[8], yielding 18,960 genes. After removal of genes that did not match with our motif scan, the COPDGene data contained 17,253 genes.

### LGRC

Gene expression data from 581 lung tissue samples in the LGRC (GSE47460) [1] was profiled using two array platforms: Agilent-014850 Whole Human Genome Microarray 4x44K G4112F and Agilent-028004 SurePrint G3 Human GE 8x60K arrays. LIMMA was used to background correct and normalize gene expression across samples within each of these two platforms. Genes that were represented by more than one probe were then removed and the expression data was merged between the two array platforms by matching probes that represented the same gene, leaving 17,573 genes. Next, batch effect due to the array platform was addressed by running ComBat [27]. Genes not present in our motif scan were then removed, yielding 14,721 genes. After normalization we filtered the samples included in the LGRC dataset by removing those that corresponded to subjects that (1) were not designated as either a COPD case or control (mostly subjects with Interstitial Lung Disease), (2) had a diagnosis of COPD, but spirometric measures in the normal range, (3) had been identified as non-Caucasian, (4) had been labeled as a former smoker, but had zero or unknown pack years, (5) had high pre-bronchodilator FEV1/FVC ratios, or (6) had been taken as a biological replicate of another sample which was included. After removal of those samples we were left with 164 COPD cases and 64 controls for which we had gene expression data.

### LTCDNM

Gene expression data from the LTCDNM (GSE76925)[39] was collected using HumanHT-12 BeadChips. Quality control was performed using quantile, signal-to-noise, correlation matrix, MA, and principal component analysis (PCA) plots using R statistical software (v 3.2.0) to identify outliers and samples with questionable or low-quality levels, distributions, or associations. This process yielded 151 samples for analysis, including 115 subjects classified as either diagnosed with COPD (87) or as a smoker control (28). After filtering for low variance and percentage of high detection p-values, 32,831 probes representing 20,794 genes were retained. The R package lumi [10] was then used for background correction, log2 transformation and quantile normalization. Finally, we collapsed probes to gene symbols based on maximum gene expression and removed genes that were not matched with our motif scan, yielding 14,273 genes.

### TFs included in analysis

For each study, we identified transcription factors for which we had gene expression data, removing those transcription factors that lacked expression values. This mapping and filtering left 164 transcription factors in ECLIPSE and COPDGene, 148 in LGRC, and 145 in LTCDNM. MONSTER was run separately on each of these studies. Comparisons of differential transcription factor involvement across studies were performed using the 143 transcription factors that were common to all four studies.

## MONSTER: MOdeling Network State Transitions from Expression and Regulatory data

The MONSTER algorithm conceptually consists of three parts: (1) inferring a gene regulatory network, (2) computing a transition matrix, and (3) quantifying the differential transcription factor involvement. We review each of these steps separately below.

### Inferring Gene Regulatory Networks

In 2013, we described PANDA [18], a method for estimating gene regulatory networks that uses “message passing” [15] to integrate multiple types of genomic data. PANDA begins with a prior regulatory network based on mapping transcription factor motifs to a reference genome and then integrates other sources of data, such as protein-protein interaction and gene expression, to estimate a collective network. While PANDA has proven to be very useful in a number of applications [31, 20, 19], its iterative approach to edge weight optimization limits its utility in situations requiring a large number of network bootstrap estimations, including applications where the sample size is large [48].

To overcome this limitation in MONSTER we developed a regression-based approach that considers the available evidence of a gene regulatory “edge” in the network for each possible transcription factor-gene pair. This evidence can be divided into two components, referred to here as direct and indirect. Consider the edge between a gene that codes for a transcription factor, *TF_i_*, and another gene. The direct evidence, *d_i,j_*, can be estimated by the squared conditional correlation:

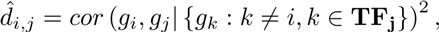

where *g_i_* is the gene which encodes *TF_i_*, *g_j_* is any other gene in the genome, and **TF_j_** is the set of gene indices corresponding to known transcription factors with binding site in the promoter region of *g_j_*. The correlation is conditioned on the expression of all other potential regulators of *g_j_* based on the transcription factor motifs associated with *g_j_*.

Naturally, the use of direct evidence alone inadequately captures regulatory relationships, which can be difficult to estimate due to systematic and technical noise as well as biological factors, such as transient protein-protein interactions and post-translational modifications, that may mask or modify a true regulatory effect. Therefore we want to complement our estimate of the likelihood of a regulatory mechanism by aggregating the information from the gene expression patterns of all suspected targets of any given transcription factor.

PANDA achieves its superior performance in part by convergence towards an “agreement” across multiple sources of evidence, in essence requiring that large collections of gene expression patterns must agree with the proposed regulatory structure in order to claim an interaction. In MONSTER, we look for agreement between the gene expression patterns of large sets of co-targeted genes. We refer to this as “indirect evidence” and estimate this by once again using the regulatory prior. Here, we no longer consider transcription factors to be members of the set of genes and instead consider each of the *m* transcription factors to be binary classifications across the entire gene list. Class labels are determined by the presence or absence of a sequence binding motif for a given transcription factor in the promoter region of a gene. For each transcription factor, we use the gene expression patterns of all targeted genes against all non-targeted genes to build a classifier. In this manner we are assigning a higher score for edges connecting each transcription factor to genes which demonstrate an expression pattern more similar to the suspected targets.

Based on this, the indirect evidence between the two nodes, *θ_i,j_*, is estimated by the fitted probability that *g_j_* belongs to the class of genes targeted by *TF_i_*. We use a logistic regression on the gene expression data with outcome taken to be the existence or non-existence of a known sequence motif for *TF_i_* in the promoter region of *g_j_*.

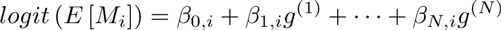

where the response *M_i_* is a binary vector of length *p* indicating the of the presence of a sequence motif for transcription factor *i* in the vicinity of each of the *p* genes. And where *g*^(*k*)^ is a vector of length *p* representing the expression of genes in sample *k*.

For a given transcription factor-gene pair, the fitted values for each *TF_i_ − g_j_* pair define the “indirect” evidence *θ_i,j_*, which can be estimated by:

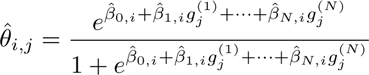

where 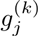 is the measured gene expression for sample *k* at gene *j*.

We score each gene according to the strength of indirect evidence for a regulatory response to each of the transcription factors and combine this with the direct evidence of regulation. Combining our measures of direct and indirect evidence presents some challenges. Though both are bounded by [0,1] their interpretations are quite different. The direct evidence can be considered in terms of its conditional gene expression *R*^2^ between nodes, while the indirect evidence is interpreted as an estimated probability. Therefore, we use a non-parametric approach to combine evidence. Specifically, the targets of each transcription factor are ranked and combined as a weighted sum, *w_i,j_* = (1 − *α*)[*rank (d*̂_*i,j*_)] + *α* [rank (*θ̂_i,j_*)], where *α* is a constant bounded between [0, 1]. Our choice of the weight is by default *α* = 0.5, corresponding to an equal contribution of direct and indirect evidence. This parameter could be adjusted if the context of a study involved reason to prefer one source of evidence over the other.

### Computation of MONSTER's transition matrix

The hypothesis behind MONSTER is that different phenotypes are characterized by distinct regulatory networks and that transitions between networks are associated with large-scale changes in the regulatory structure of the network. Essentially, transcription factors gain or lose targets and in doing so, alter the structure of the network from one phenotypic state to another. The task of identifying meaningful network transitions then becomes an evaluation of the relative refinement of edge weights.

Our analysis of validation data sets (shown below) indicates that the reconstructed networks are strongly driven by the structure of the motif prior, with small changes defining differences between phenotypes. Hence, in comparing networks between phenotypes, the problem becomes one of of understanding changes in edges that have relatively low signal and high noise. In other words, state transitions are characterized by a large number of individually unreliable edge weights.

Consider two adjacency matrices, **A** and **B**, that represent two gene regulatory networks estimated from a case-control study. Each matrix has dimensions (*p* × *m*) representing the set of *p* genes targeted by *m* transcriptionfactors. We seek a matrix, **T**, such that

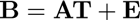

where **E** is our error matrix, which we want to minimize. Intuitively, we may frame this as a set of *m* independent regression problems, where *m* is the number of transcription factors and also the column rank of **A**, **B**, **T**, and **E**. For a column in **B**, **b***_i_*, we note that a corresponding column in **T**, *τ_i_*, represents the ordinary least squares solution to

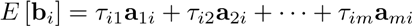

or alternativelyexpressed

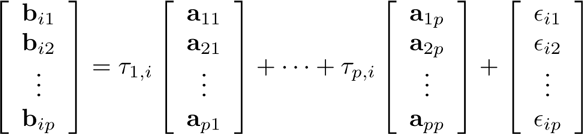

where *E* [*ϵ_ij_*] = 0. This can be solved with normal equations,

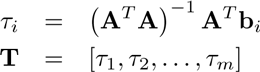

which produces the least squares estimate. In other words, the loss function 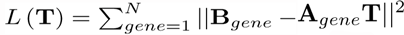 is minimized.

It is easy to see how this allows for a straightforward extension via the inclusion of a penalty term. For example, an *L*_1_ regularization[47] can be used to create an identity penalty model matrix for each column regression such that only the *k^th^* diagonal element is 0 and all other diagonals are 1. This gives unpenalized priority for the *k^th^* regression coefficient in the *k^th^* regression model:

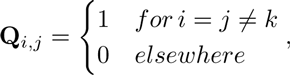

which results in the minimization of the penalized residual sum of squares

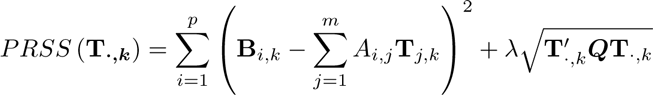

Although not used in the analysis presented in the main text, an implementation of this extension is available in the R package MONSTER.

### Analyzing the Transition Matrix

The derivation described above illustrates a key feature of the MONSTER method. Specifically, that the transition matrix (**T**) reduces the case-control network transformation from a set of 2 × *p* × *m* estimates to a set of *m* × *m* estimates that are more easily interpreted. We can think of a column, *τ_i_*, on the matrix **T** as containing the linear combination of regulatory targets of *TF_i_* in **A** that best approximates the regulatory targets of *TF_i_* in **B**. As one would expect, a large proportion of the matrix “mass” would be on the diagonal for those transcription factors which do not change regulatory behavior between case and control. It is therefore of interest to evaluate values off of the diagonal as indicative of a network transition.

There are many biological processes involved in gene regulation that may differ between phenotypic states, including RNA degradation, post-translational modification, protein-level interactions and epigenetic alterations. These all have the ability to impact transcription factor targeting without impacting the expression level of the transcription factor itself. Because our hypothesis is that changes in phenotype are associated with changes in regulatory networks, we want to identify those transcription factors that have undergone significant overall changes in behavior between states. As a measure to quantify such changes, we define the differential Transcription Factor Involvement (dTFI),

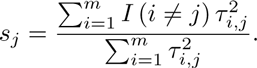

The dTFI can be loosely interpreted as the proportion of transcription factor targeting that is gained from or lost to other available transcription factors as the state changes. It is a statistic on the interval [0, 1] that can be used to identify transitions which are systematic, informative, and non-arbitrary in nature. In other words, the dTFI can capture edge weight signal for which there is an attributable regulatory pattern based on the inferred networks.

**Supporting Figure 1:**
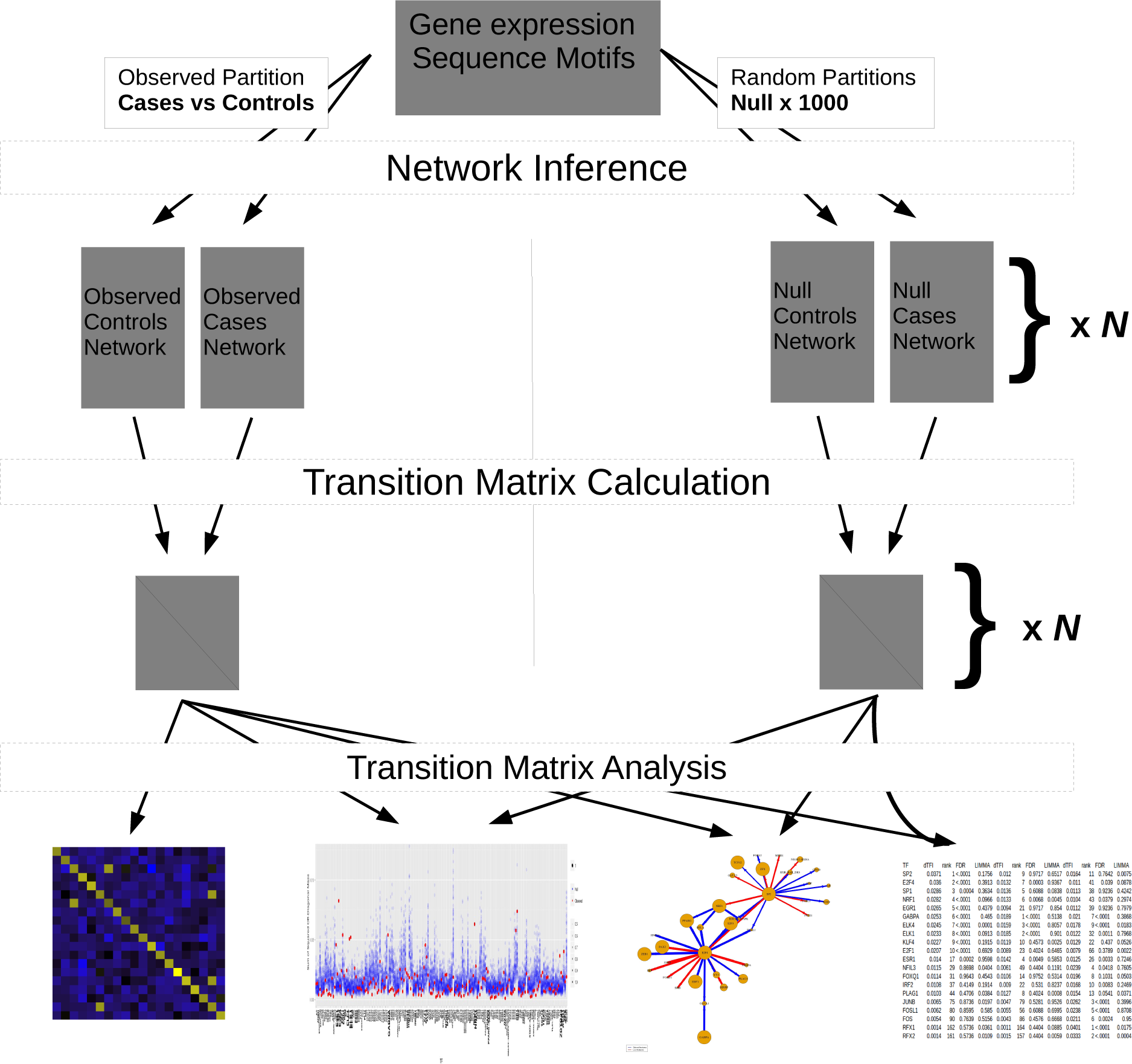
Overview of MONSTER analysis workflow. (1) Network inference is computed separately to subsets of the gene expression data including the case group, the control group and *N* permutations of the case and control labels. (2) The transition matrix is estimated between the cases and controls and each of the pairs of permuted “case” and “control” groups. (3) The transition matrix computed between the case and control group is interpreted within the context of the *N* matrices estimated for the permuted groups.

The distribution of the dTFI statistic under the null has a mean and standard deviation that depends to a large extent on the motif-based network prior structure. In particular, we find that both mean and standard deviation of the dTFI are higher for transcription factors that have fewer prior regulatory targets. From a statistical perspective, transcription factors with relatively more targets are able to generate more stable targeted expression patterns, which leads to more consistent estimates in “agreement”. From a biological perspective, increased motif presence may indicate that transcription factors are more likely to be involved in “housekeeping” or tissue specific processes that are unlikely to change between cases and controls.

We address the dependence of the null distribution of the dTFI on the motif structure using the following resampling procedure (Supporting Figure 1):

0. Gene regulatory networks are reconstructed based on a prior regulatory structure and gene expression from case and control samples and the transition matrix and the dTFI values for each transcription factor are computed.
1. Gene expression samples are randomly assigned as case and control forming null-case and null-control groups with sizes reflecting the true case and control groups.
2. Gene regulatory networks are reconstructed for the null-case and null-control groups with the same prior regulatory structure.
3. The transition matrix algorithm is applied to the two null networks.
4. The dTFI is calculated for each transcription factor based on the computed null transition matrix.
5. Steps 1-4 are repeated *n* times.

For the analysis presented in the main text, we set *n* = 400. This procedure allows us to estimate a background distribution of dTFI values based on the underlying motif prior network structure and therefore test the significance of observed dTFI values between cases and controls.

## Validation of the MONSTER Approach

### MONSTER recovers network edges in in silico, Escherichia coli and Yeast (Saccharomyces cerevisiae)

For its initial step, MONSTER uses gene expression together with a prior network structure to infer regulatory network edges. For method testing and validation of MONSTER's network estimates we used four data sets of increasing biological complexity: (1) in silico, (2) Escherichia coli, and (3) Saccharomyces cerevisiae (yeast) expression data together with simulated motif priors derived from reference networks and (4) yeast expression data together with a biological motif prior generated independently of the reference. For data set (4), we used the yeast motif prior, 106 gene expression samples from transcription factor knockout or overexpression conditions, and ChIP gold standard described in Glass et. al.[18]. Data for the first three sources was obtained from the 2012 DREAM5 challenge data set[33]. This challenge asked contestants to infer gene networks from expression data alone, using a reference standard for evaluation. For the purposes of validating MONSTER, we instead started with the reference network and randomly perturbed TF-gene pairs to create the type I and type II error rates consistent with biological yeast motif prior used in the fourth data set. Specifically, if an edge appeared in the reference network, that edge appeared in the simulated motif data with probability 0.3; if an edge was absent from the reference network, that edge appeared in the simulated motif data with probability 0.1. These probabilities result in an area under the Receiver-Operator Characteristic curve (AUC-ROC) of approximately 0.7 for prediction of the reference edges by the simulated edges.

For each of the data sets, we evaluated the accuracy of MONSTER's network inference method using AUC-ROC. For the DREAM5 data sets we applied MONSTER to the expression data together with the simulated priors and used the original reference networks as our gold-standards. For the fourth data set we applied MONSTER to the expression and motif data, and used the ChIP-chip data as our gold-standard. We found that in all four of these data sets, the accuracy of the estimated edges from MONSTER's network inference was superior to the accuracy of the input motif prior data (Supporting Figure 2).

**Supporting Table 1:** Comparison of edge weight difference to Transition Matrix in simulated case-control gene expression. Several network inference methods were run on our in silico case-control data. The overall network area under the curve of the receiver-operator characteristic (AUC-ROC) was performed for each method averaged across cases and controls. The naive transcription factor-transcription factor transitions were calculated as the difference in transcription factor-transcription factor edge weight between cases and controls. The transition matrix transcription factor-transcription factor transitions used the absolute transition matrix values.

| transcription factor | ECLIPSE |  |  | COPDGene |  |  | LGRC |  |  | LTCDNM |  |  |
| --- | --- | --- | --- | --- | --- | --- | --- | --- | --- | --- | --- | --- |
|  | dTFI | rank | FDR | dTFI | rank | FDR | dTFI | rank | FDR | dTFI | rank | FDR |
| SP2 | .0314 | 1 | .0357 | .0100 | 9 | .6812 | .0213 | 6 | .3752 | .0176 | 2 | .7438 |
| E2F4 | .0236 | 2 | <.0001 | .0143 | 3 | <.0001 | .0160 | 14 | .037 | .0148 | 7 | <.0001 |
| SP1 | .0230 | 3 | .1551 | .0089 | 18 | .7721 | .0179 | 10 | .3594 | .0169 | 4 | .5516 |
| ZNF263 | .0226 | 4 | .311 | .0089 | 16 | .3372 | .0177 | 11 | .7716 | .0152 | 6 | .927 |
| EGR1 | .0224 | 5 | .1242 | .0079 | 23 | .7597 | .0124 | 28 | .6892 | .0152 | 5 | .5305 |
| NRF1 | .0196 | 6 | <.0001 | .0115 | 5 | .0304 | .0122 | 30 | <.0001 | .0139 | 11 | .0558 |
| GABPA | .0185 | 7 | <.0001 | .0157 | 2 | <.0001 | .0176 | 12 | <.0001 | .0097 | 32 | .0853 |
| ELK1 | .0177 | 8 | <.0001 | .0174 | 1 | <.0001 | .0151 | 17 | <.0001 | .0083 | 40 | .2099 |
| ZFX | .0175 | 9 | <.0001 | .0076 | 24 | .8366 | .0103 | 40 | .4348 | .0132 | 16 | .2739 |
| KLF4 | .0173 | 10 | .1025 | .0072 | 28 | .8142 | .0143 | 21 | .2312 | .0119 | 20 | .5516 |
| ESR1 | .0169 | 11 | .0357 | .0106 | 7 | .0941 | .0127 | 27 | .0888 | .0176 | 3 | <.0001 |
| ELK4 | .0168 | 12 | <.0001 | .0125 | 4 | <.0001 | .0152 | 16 | <.0001 | .0086 | 39 | .1318 |
| TFAP2C | .0139 | 17 | .0656 | .0114 | 6 | .0941 | .0148 | 19 | .037 | .0121 | 19 | .2099 |
| PLAG1 | .0124 | 21 | .263 | .0092 | 15 | .4136 | .0219 | 5 | <.0001 | .0146 | 8 | .1554 |
| FOXQ1 | .0115 | 28 | .9318 | .0099 | 10 | .7905 | .0209 | 7 | .2846 | .0107 | 27 | .927 |
| FOSL1 | .0082 | 57 | .9175 | .0061 | 41 | .6166 | .0220 | 4 | .037 | .0131 | 17 | .3496 |
| NFIL3 | .0077 | 62 | .2365 | .0067 | 33 | .0304 | .0264 | 1 | .4669 | .0209 | 1 | .7121 |
| FOS | .0068 | 73 | .9175 | .0057 | 48 | .5212 | .0198 | 9 | .037 | .0112 | 24 | .5139 |
| JUNB | .0067 | 77 | .9318 | .0059 | 43 | .6392 | .0236 | 2 | <.0001 | .0146 | 9 | .2299 |
| RFX1 | .0019 | 159 | .3532 | .0009 | 164 | <.0001 | .0233 | 3 | <.0001 | .0070 | 48 | .3496 |
| RFX2 | .0019 | 158 | .4041 | .0012 | 163 | .0482 | .0200 | 8 | <.0001 | .0049 | 81 | .6245 |

Additional Table 1: **Top Transcription Factor Hits.**
| transcription factor | ECLIPSE |  | COPDGene |  | LGRC |  | LTCDNM |  |
| --- | --- | --- | --- | --- | --- | --- | --- | --- |
|  | dTFI rank | LIMMA p | dTFI rank | LIMMA p | dTFI rank | LIMMA p | dTFI rank | LIMMA p |
| SP2 | 1 | .1756 | 9 | .6517 | 6 | .0075 | 2 | .0009 |
| E2F4 | 2 | .3913 | 3 | .9367 | 14 | .0878 | 7 | .8232 |
| SP1 | 3 | .3634 | 18 | .0838 | 10 | .4242 | 4 | .9759 |
| ZNF263 | 4 | .9834 | 16 | .0028 | 11 | .0271 | 6 | .1859 |
| EGR1 | 5 | .4379 | 23 | .8540 | 28 | .7979 | 5 | .0378 |
| NRF1 | 6 | .0966 | 5 | .0045 | 30 | .2974 | 11 | .3418 |
| GABPA | 7 | .4650 | 2 | .5138 | 12 | .3868 | 32 | .5771 |
| ELK1 | 8 | .0913 | 1 | .9010 | 17 | .7968 | 40 | .0005 |
| ZFX | 9 | .8253 | 24 | .5795 | 40 | .0474 | 16 | .1572 |
| KLF4 | 10 | .1915 | 28 | .0025 | 21 | .0526 | 20 | .1159 |
| ESR1 | 11 | .9598 | 7 | .5853 | 27 | .7246 | 3 | .3477 |
| ELK4 | 12 | .0001 | 4 | .8057 | 16 | .0183 | 39 | .7314 |
| TFAP2C | 17 | .2318 | 6 | .9574 | 19 | .5853 | 19 | .6754 |
| PLAG1 | 21 | .0384 | 15 | .0008 | 5 | .0371 | 8 | .9523 |
| FOXQ1 | 28 | .4543 | 10 | .5314 | 7 | .0503 | 27 | .5340 |
| FOSL1 | 57 | .5850 | 41 | .6995 | 4 | .8708 | 17 | .3686 |
| NFIL3 | 62 | .0404 | 33 | .1191 | 1 | .7605 | 1 | .8650 |
| FOS | 73 | .5156 | 48 | .6668 | 9 | .9500 | 24 | .7891 |
| JUNB | 77 | .0197 | 43 | .9526 | 2 | .3996 | 9 | .6077 |
| RFX1 | 159 | .0361 | 164 | .0885 | 3 | .0175 | 48 | .8285 |
| RFX2 | 158 | .0109 | 163 | .0059 | 8 | .0004 | 81 | .1345 |

### MONSTER accurately predicts transcription factor transitions in in silico gene expression data

We next used simulated data to evaluate MONSTER's transition matrix. To begin, we randomly generated a “true” control adjacency matrix, **M**_0_, which contained information for all possible edges between *m* = 100 transcription factors and *p* = 10, 000 genes with “edge weights” sampled from a standard uniform distribution. We then defined a state transition matrix, T, with diagonal elements set equal to one and 1, 000 random off-diagonal elements (representing random pairs of transcription factors) set equal to values sampled from a uniform random distribution between −1.0 and 1.0. These off-diagonal elements (transcription factor pairs) ultimately represent the transitions that we seek to recover and their corresponding values represent the magnitude of the regulatory transition. Finally, based on **M**_0_ and T we defined the “true” cases network as **M**_1_ = T**M**_0_.

Next, we generated two in silico gene expression datasets, one each for the case and control networks. To do this, we sampled 500 times from each of two multivariate Gaussian distributions with the variance-covariance matrix, Σ, defined as 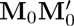 and 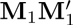 for controls and cases, respectively. We note that we scaled the magnitude of the diagonal elements of Σ by 4 to simulate noise in the in silico data. This value was chosen such that the networks predicted using the in silico gene expression data had an AUC-ROC of approximately .70 when evaluated using the “true” networks (see below).

We next used this simulated data to reconstruct networks using several commonly used network inference methods, including the Pearson correlation (used in WGCNA) [29] [30], Topological Overlap Measure (TOM) [41], Algorithm for the Reconstruction of Gene Regulatory Networks (ARACNE) [34], and Context Likelihood of Relatedness (CLR) [13]. The implementation of each method was from the R package nettools [14].

We next constructed a gold-standard for our network transitions, defined as **T***_GS_* = ceil([**M**]). For each of the five network inference methods, we then evaluated the accuracy of two potential approaches for identifying network alterations. First, we simply subtracted edge weights between the inferred cases network and the inferred controls network and selected those edges that extended between the 100 TFs in our model (excluding those genes that were not TFs). Second, we used MONSTER to predict the transition needed to map the control network to the case network. The results are summarized in Supporting Table 1. For each of the network inference methods tested, we found that the transition matrix showed substantial improvement over the edge weight difference method in identifying transitions between transcription factors. In all cases, the edge weight difference (column 3) was not statistically significant for predicting transitions, but when the transition matrix was used (column 4) a strong predictive signal appeared.

**Supporting Figure 2:**
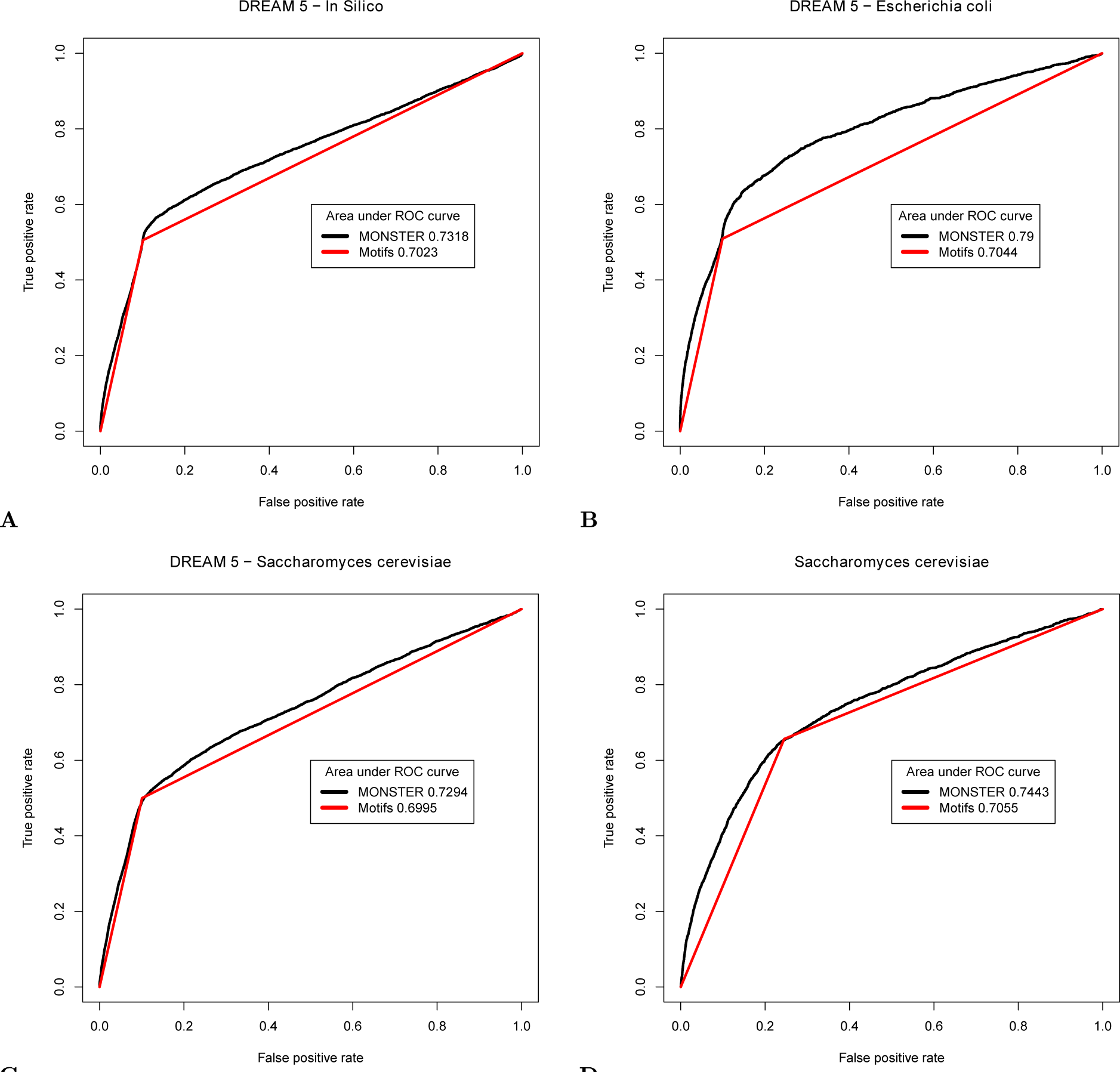
Receiver-Operator Characteristic curves for three DREAM 5 data sets (A) in silico, (B) Escherichia coli, (C) Saccharomyces cerevisiae, and an (D) additional Saccharomyces cerevisiae data set as described in Glass et. al.[18]. The prior network for each of the DREAM5 data set analyses was derived from the validation standard, with error introduced (both type I and type II) bringing the area under the ROC curve to ≈ 0.70. In the other Saccharomyces cerevisiae data set analysis, sequence motifs were used as the prior and a ChIP-chip derived network was used as the validation standard. In each of these tests, we observed a measurable improvement in performance of MONSTER's network inference method over the prior.

### MONSTER finds significant protein-protein interactions

There are numerous biological regulatory mechanisms that may play a role in transitions between phenotypic states. Of particular interest to us are those that are not readily detectable via conventional methods for the analysis of gene expression data. For example, gene regulation involves complex processes in which transcription factors, either singly or in multiprotein complexes, bind to DNA in the region of a gene to activate or repress the transcriptional process. Such multi-protein interactions create combinatorial complexity that can explain much of the variation in organism complexity which is unexplained by gene expression alone [32].

As reported in the main text, we ran MONSTER on data from 84 smoker controls and 136 COPD subjects in the ECLIPSE study. To test whether MONSTER could reliably detect protein-protein interactions between regulatory transcription factors, we evaluated whether our estimated transitions between case and control COPD networks in this analysis recapitulated known protein-protein interactions, as reported in Ravasi et. al.[40] and processed in Glass et. al.[20]. This dataset contained 223 interactions between the transcription factors we used as input of our model; of these, 39 were self-interacting and were removed. We attempted to predict the remaining 184 interactions between transcription factors using MONSTER.

We used the absolute value of the transition matrix and tested whether that value predicted protein-protein interactions based on the area under the ROC curve. To assess the significance of AUC-ROC, we also applied this evaluation to the 400 “random” transition matrices generated based on the randomized phenotypic labels. MONSTER achieved an AUC-ROC score of .548, suggesting predictive power to identify known PPI between transcription factors. While weak, this result exceeded all randomized phenotype results and was significant at *p* < .0025. This indicates that MONSTER is able to extract a small but significant protein interaction signal from highly obfuscated data.

### Irreproducibility of network inference methods in estimating transcription factor - gene edge-weights in COPD

Conceptually, MONSTER is comprised of two elements. The first infers gene regulatory networks from transcriptional data while the second uses the networks inferred for two different phenotypes to calculate the transition matrix between states. Instead of using the second part of the MONSTER approach to understand the transition between one state and another, one could imagine instead substracting the edge-weights predicted for two networks and using those differences to define a transition between two phenotypic states. To test whether this is a reasonable approach we examined the reproducibility of edge weight differences between case and control networks estimated for four COPD datasets using MONSTER's network reconstruction approach as well as three other widely used network inference methods: Algorithm for the Reconstruction of Gene Regulatory Networks (ARACNE), Context Likelihood of Relatedness (CLR), and the standard Pearson correlation used in such methods as Weighted Gene Correlation Network Analysis (WGCNA).

We used each of the four methods to separately estimate networks for cases and controls in each of the COPD studies. We then calculated the difference between case and control edges (differential edge weights) in each study for each method. We reasoned that if edge-differences were reflective of biologically meaningful associations, these should be present in each study and should appear as a correlated set of differential edge weights.

We plotted the differential edge weights for each pairwise combination of studies (Supporting Figure 3) and found that the differential edges found by ARACNE, CLR, WGCNA and MONSTER were almost entirely study specific, meaning that edges are found in one study comparing smoker controls to COPD patients are not found in a second study comparing the same phenotypes. Clearly, evaluation of individual edge-weight differences is not a reproducible approach for comparing inferred networks and stands in stark contrast to the highly reproducible set of differentially-involved set of transcription factors that we were able to identify across all four studies (as presented in the main text).

**Additional Table 1:**
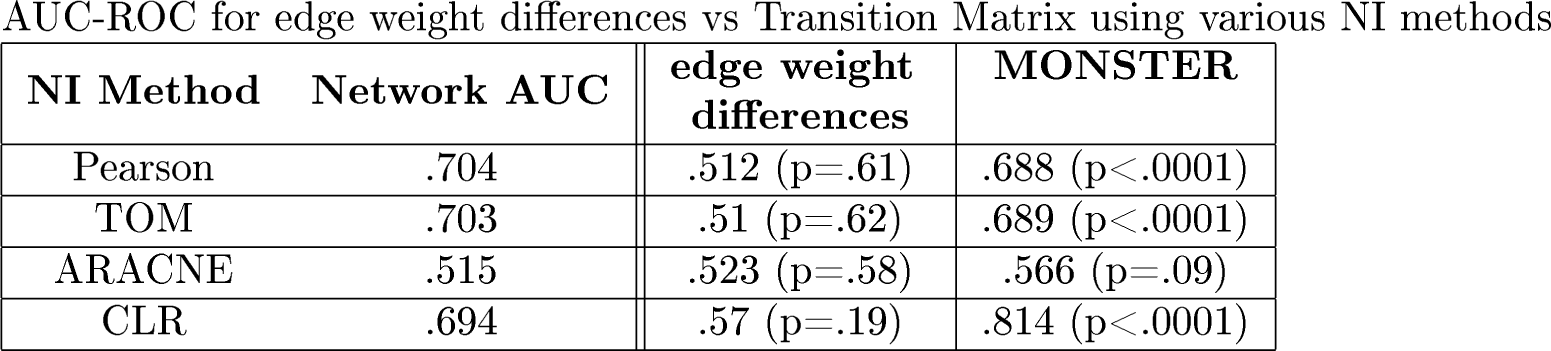
Top Transcription Factor Hits. **A** Combined list of transcription factors which were among the top 10 hits (out of 166 available transcription factors) in any of the 4 studies, ordered by the dTFI in the ECLIPSE study. For each study, columns indicate the transcription factor's (1) differential transcription factor Involvement, (2) dTFI Rank within list of transcription factors, (3) and Significance of dTFI by false discovery rate. **B** The same list of top transcription factors evaluated for differential gene expression analysis using LIMMA. A substantial number of differentially involved transcription factors do not exhibit gene expression differentiation, highlighting the ability of MONSTER to identify key features distinguishing phenotypes which are not detectable via gene expression analysis.

**Supporting Figure 3:**
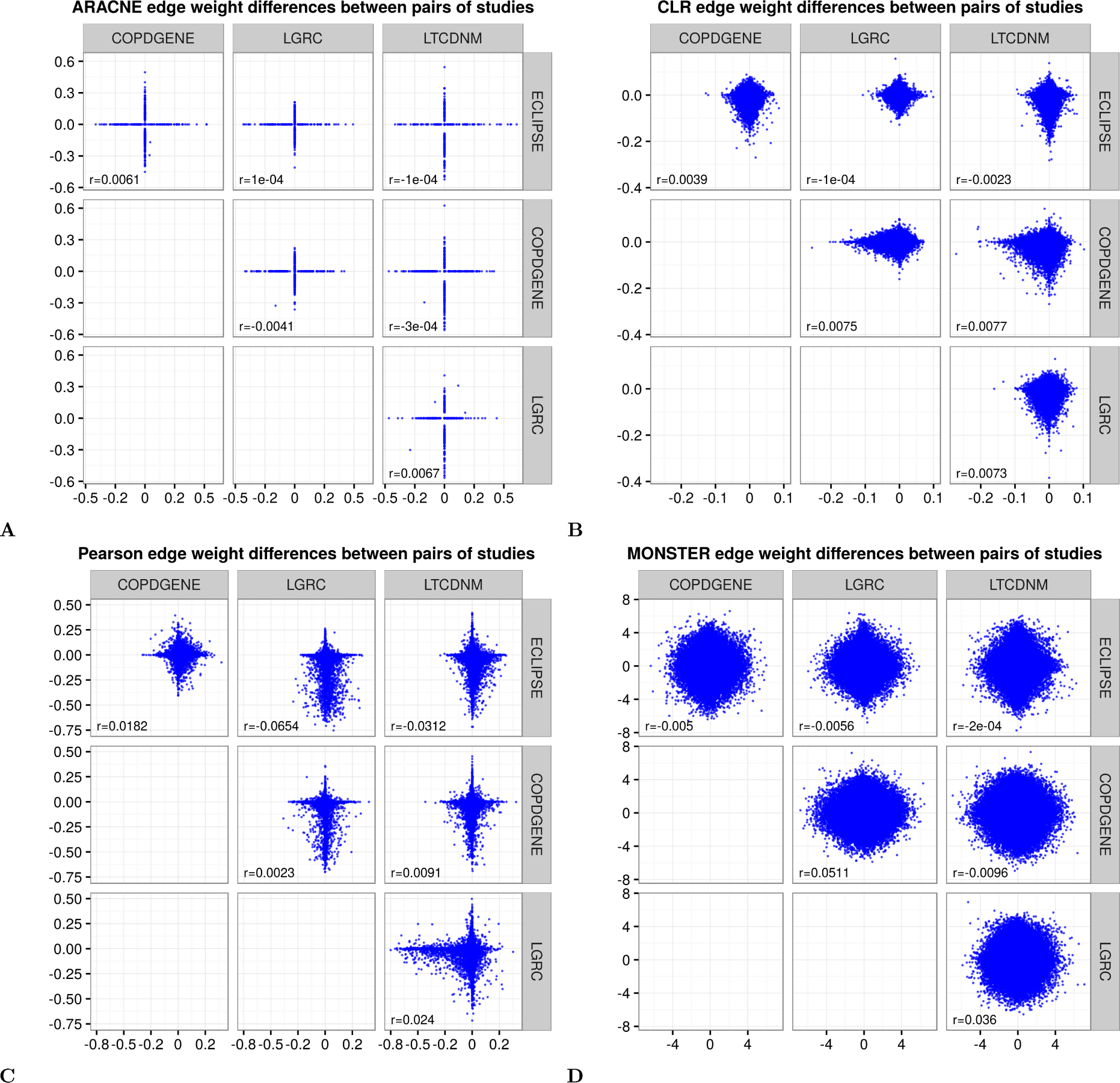
Edge weight differences between cases and controls do not correlate across studies. Using MONSTER and three other commonly used methods, we performed network inference separately on cases and controls in four COPD data sets. Here, the case-control difference is compared for each method in each data set. Most methods had very poor overall concordance in the edge weight differences they estimated. The methods tested were **A** Algorithm for the Reconstruction of Gene Regulatory Networks (ARACNE), **B** Context Likelihood of Relatedness (CLR), **C** Pearson correlation networks, such as in Weighted Gene Correlation Network Analysis (WGCNA), and **D** MONSTER. No detectable agreement between studies exist were found, regardless of network inference method or tissue type.

**Additional Figure 1:**
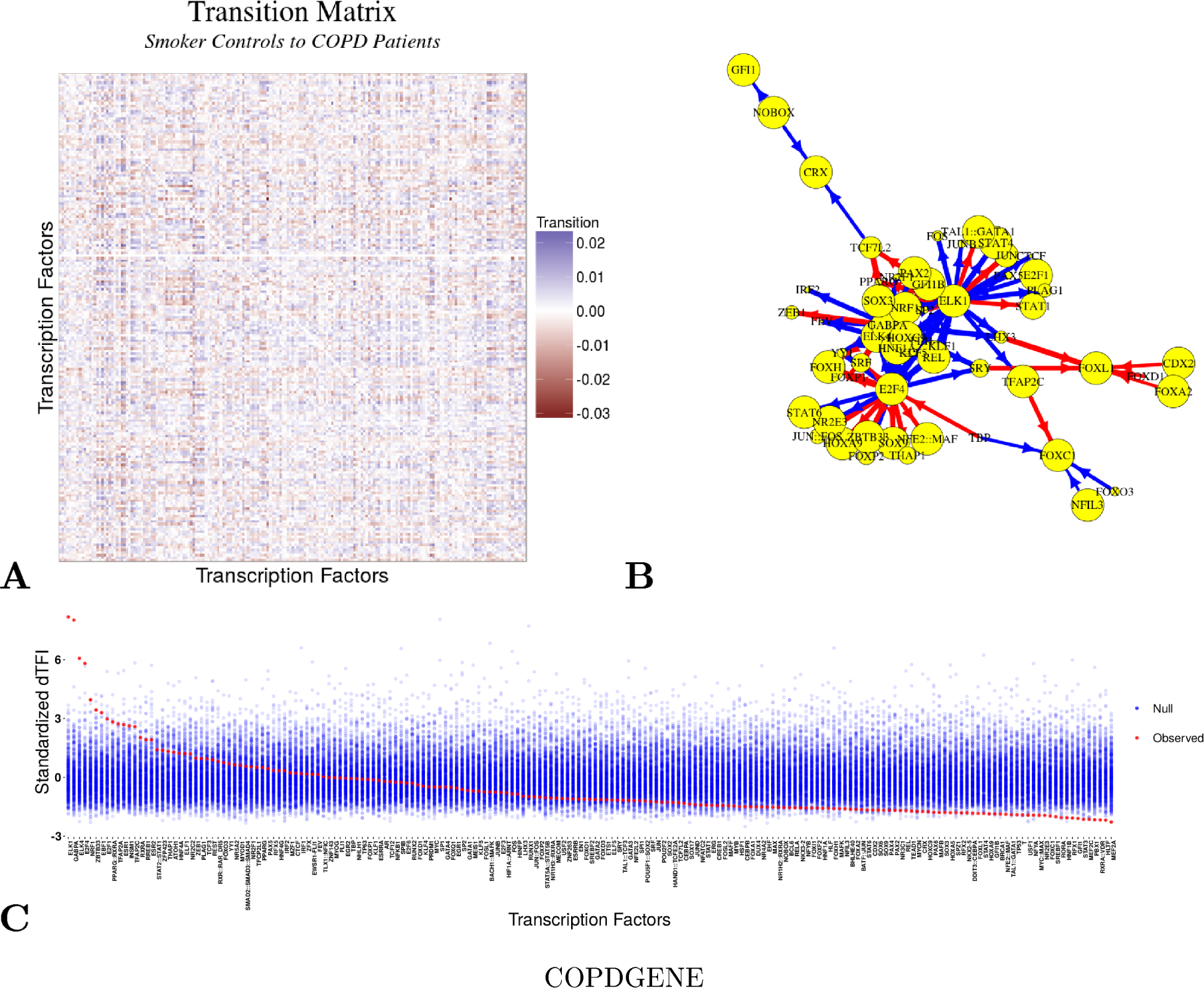
MONSTER analysis results for COPDGENE study. **A** Heatmap depicting the transition matrix calculated from smoker controls to COPD cases by applying MONSTER to the COPDGene study. For the purposes of visualization, the magnitude of the diagonal is set to zero. **B** A network visualization of the strongest 100 transitions identified based on the transition matrix shown in **A**. Arrows indicate a change in edges from a transcription factor in the Control network to resemble those of a transcription factor in the COPD network. Edges are sized according to the magnitude of the transition and nodes (transcription factors) are sized by the dTFI for that transcription factor. The gain of targeting features is indicated by the color blue while the loss of features is indicated by red. **C** The dTFI score from MONSTER (red) and the background null distribution of dTFI values (blue) as estimated by 400 random sample permutations of the data.

**Additional Figure 2:**
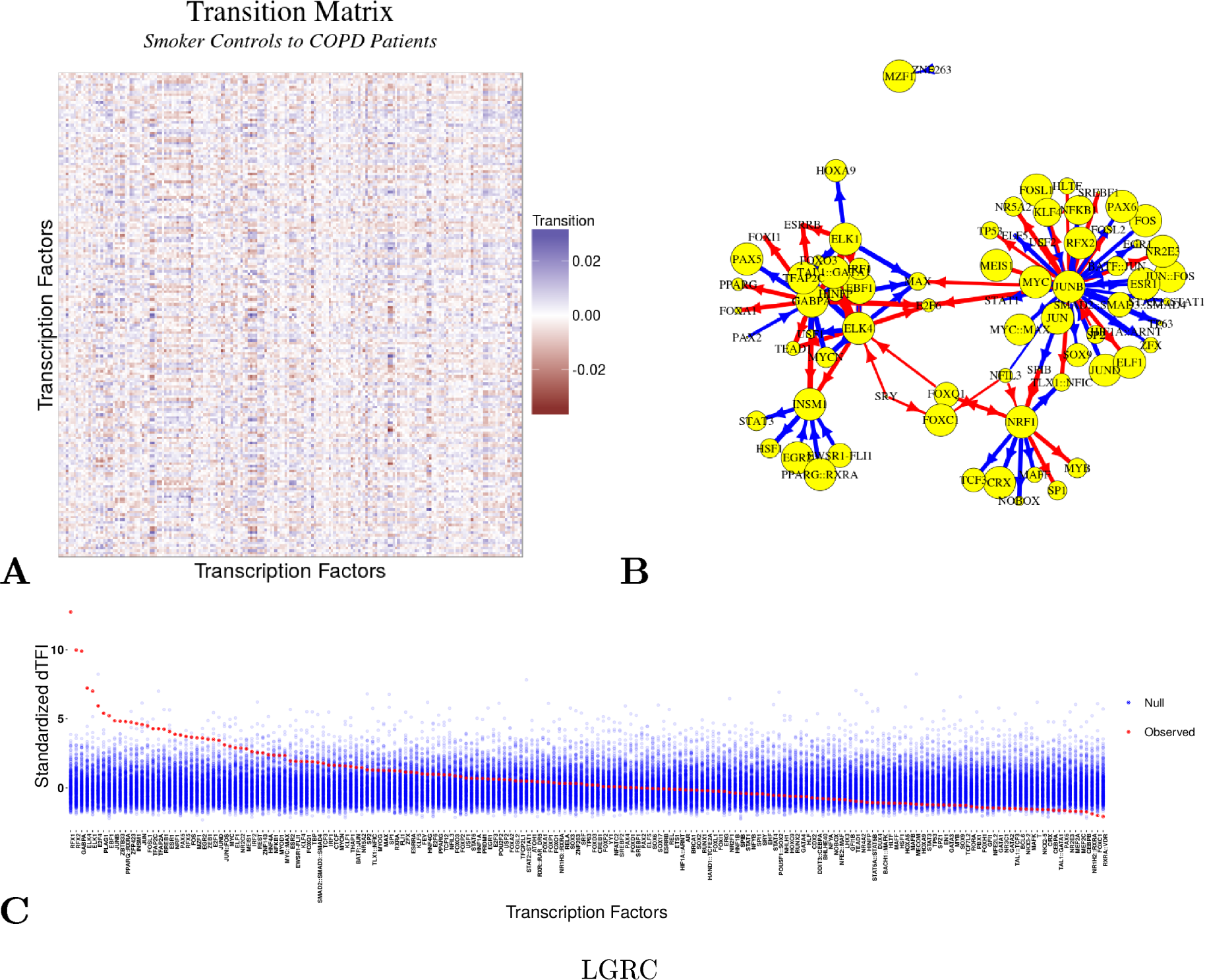
MONSTER analysis results for LGRC study. **A** Heatmap depicting the transition matrix calculated from smoker controls to COPD cases by applying MONSTER to the LGRC study. For the purposes of visualization, the magnitude of the diagonal is set to zero. **B** A network visualization of the strongest 100 transitions identified based on the transition matrix shown in **A**. Arrows indicate a change in edges from a transcription factor in the Control network to resemble those of a transcription factor in the COPD network. Edges are sized according to the magnitude of the transition and nodes (transcription factors) are sized by the dTFI for that transcription factor. The gain of targeting features is indicated by the color blue while the loss of features is indicated by red. C The dTFI score from MONSTER (red) and the background null distribution of dTFI values (blue) as estimated by 400 random sample permutations of the data.

**Additional Figure 3:**
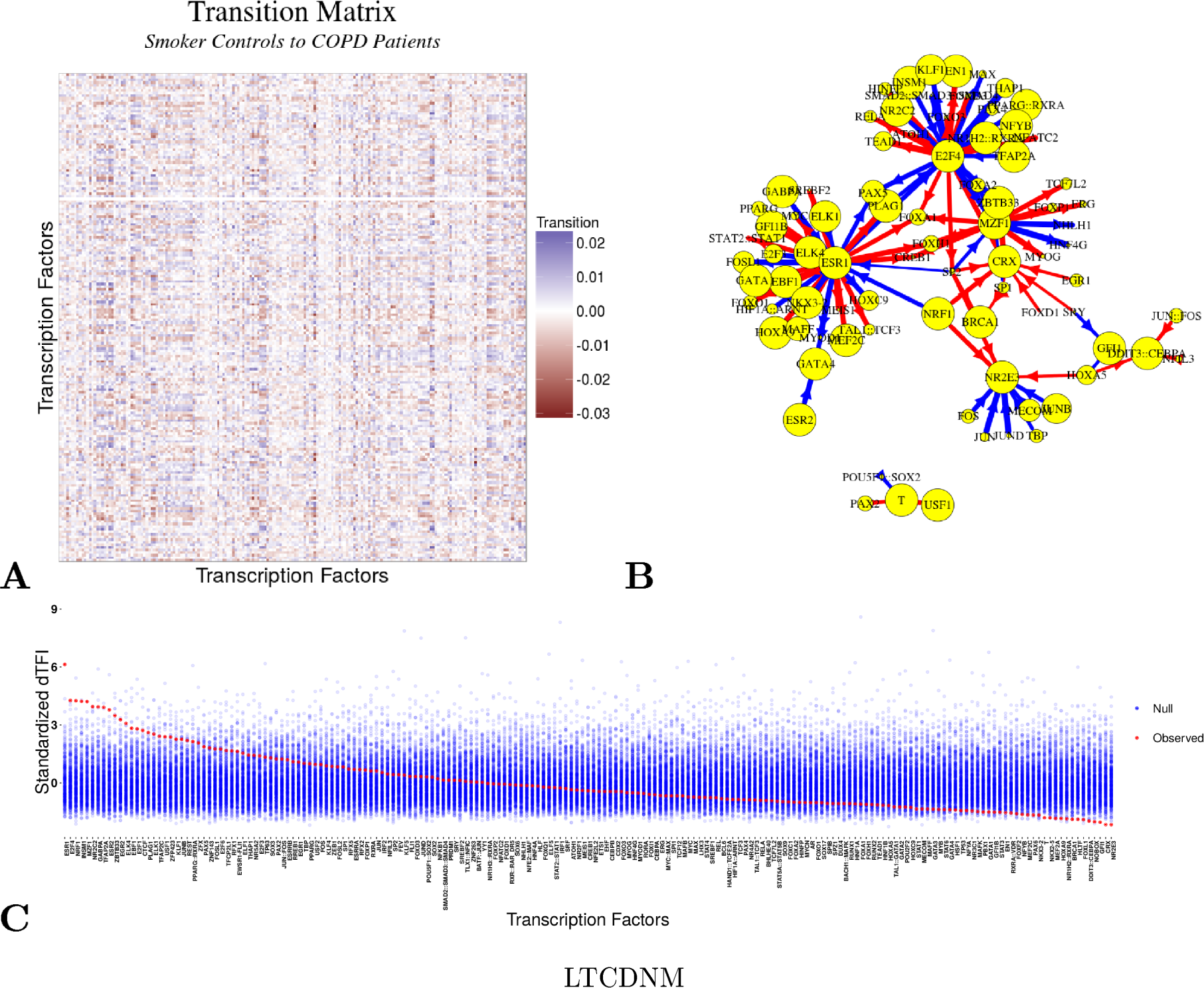
MONSTER analysis results for LTCDNM study. **A** Heatmap depicting the transition matrix calculated from smoker controls to COPD cases by applying MONSTER to the LTCDNM study. For the purposes of visualization, the magnitude of the diagonal is set to zero. **B** A network visualization of the strongest 100 transitions identified based on the transition matrix shown in **A**. Arrows indicate a change in edges from a transcription factor in the Control network to resemble those of a transcription factor in the COPD network. Edges are sized according to the magnitude of the transition and nodes (transcription factors) are sized by the dTFI for that transcription factor. The gain of targeting features is indicated by the color blue while the loss of features is indicated by red. C The dTFI score from MONSTER (red) and the background null distribution of dTFI values (blue) as estimated by 400 random sample permutations of the data.

**Additional Figure 4:**
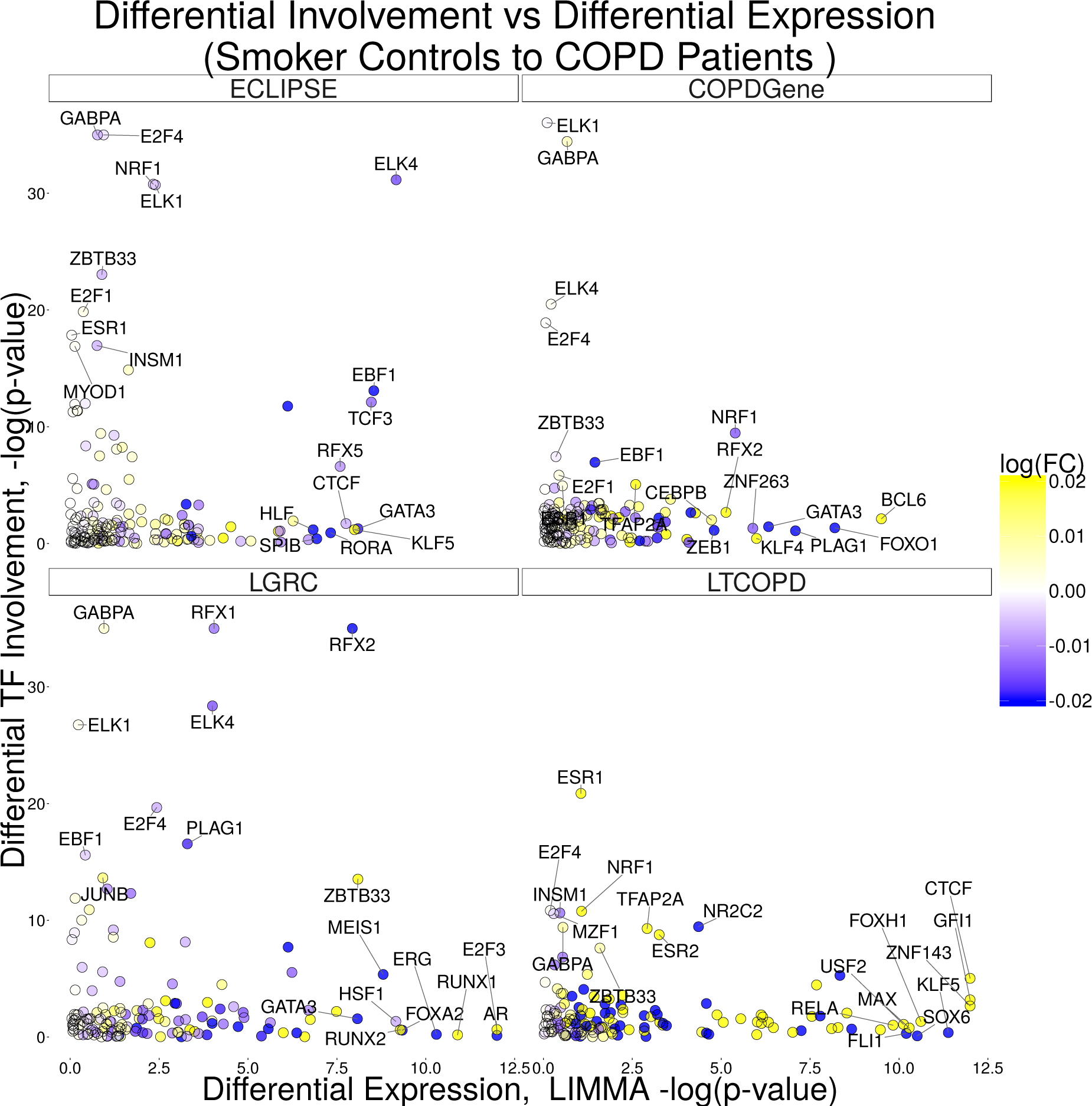
Differentially transcription factor involvement vs differential gene expression in four studies of COPD. Plots of the differential expression of transcription factors based on LIMMA, and their different involvement (dTF1) based on MONSTER. We observe much higher consistency between the transcription factors highlighted using MONSTER compared to LIMMA. In addition, we note that MONSTER commonly finds transcription factors which are differentially involved but are expressed at similar levels across cases and controls. This demonstrates the unique potential MONSTER has for discovery beyond standard gene expression analysis.

